# Adaptation to local climate in a multi-trait space: evidence from silver fir (*Abies alba* Mill.) populations across a heterogeneous environment

**DOI:** 10.1101/292540

**Authors:** Katalin Csilléry, Otso Ovaskainen, Christoph Sperisen, Alex Widmer, Felix Gugerli

**Author notes:** Corresponding author: Katalin Csilléry &. **Data archival:** SNP data will be archived on Dryad.

## Abstract

Attempts to identify and understand selection pressures responsible for local adaptation are central to evolutionary research. We tested whether populations of silver fir (*Abies alba* Mill.), sampled across a heterogeneous environment, have more strongly diverged at quantitative traits than expected from genetic drift. We genotyped 387 trees from 19 Swiss populations at 374 single-nucleotide polymorphisms (SNPs) to estimate their demographic distances, and used these to generate a null expectation for divergence in a multi-trait space including morphology and life-history traits obtained from a published common garden trial. Local soil and historical climate data were used to identify the selective environment of the source populations. Our results revealed a strong selection on height driven by temperature: trees from warm sites evolved to become taller than those from cooler sites. The evolution of growth rate, growth duration and bud break were correlated, and populations evolved towards two extreme strategies, “start early and grow slow” or “start late and grow fast”, driven by precipitation seasonality. We conclude that local climate has shaped the morphology and life-history of silver fir populations since they recolonized the Alps. Our methodology provides a show-case for empirical evaluation of adaptive evolutionary strategies combining genetic data and common gardens.

## Introduction

With increasing concerns about climate change, estimating the adaptive potential of forest trees has high societal and economic relevance. The capacity of many forest tree species to survive and reproduce across a broad geographical range is, at least partly, due to adaptation to local conditions (*e.g*. Aitken et *al*., 2008; Franks et *al*., 2014). There is growing evidence that adaptation can be important for enabling populations to persist under changing conditions, including climate change (Hoffmann & Sgrò, 2011). However, many populations are increasingly exposed to novel environmental conditions that are beyond the range to which they are adapted and/or outside of their physiological tolerance (Shaw & Etterson, 2012), and it is unclear how commonly and rapidly adaptations or plastic responses would arise in order to keep up with climate change (e.g. Franks et *al*., 2014).

Many forest tree species have distribution ranges that expand over large geographic regions, which is, at least in the Northern Hemisphere, a result of recent population expansions after the Last Glacial Maximum (Hampe & Jump, 2011). Given the large effective population sizes, predominant outcrossing mode of reproduction, and moderate to high levels of genetic diversity, forest tree species have been able to adapt to local conditions, as reflected by ample empirical evidence of trait divergence patterns that are congruent with latitudinal and altitudinal climatic gradients (Savolainen et *al*., 2007; Alberto et *al*., 2013). However, since many of these gradients also coincide with recent population expansion, it can be difficult to tease apart neutral and selective processes underlying trait variation (e.g. Yeaman et *al*., 2016).

Attempts to identify adaptive trait differentiation in forest trees have often relied on the long tradition of provenance trials, in which different populations from across the geographic range are tested in a common environment. Phenotypic data is often exploited using so-called *Q_ST_* – *F_ST_* tests, which compare quantitative trait differentiation (*Q_ST_*) with genetic divergence inferred from neutral markers (*F_ST_*) (Whitlock, 2008; Whitlock & Guillaume, 2009). A recent literature survey of forest trees suggests that most tree traits exhibit considerable heritable trait variation and that *Q_ST_* most often (69% of the studies)exceeds *F_ST_* (Lind *et al*., 2018). However, a simple comparison between *Q_ST_* and *F_ST_* has many shortcomings that have been criticized (e.g. reviewed in Leinonen et *al*. (2013)), and alternative solutions have been suggested (Chenoweth & Blows, 2008; Martin et *al*., 2008; Ovaskainen et *al*., 2011). The fundamental issue is that both *Q_ST_* and *F_ST_* are random variables, i.e. the observed values are just a single realization of the evolutionary process. Even though Whitlock & Guillaume (2009) proposed an improved *Q_ST_* – *F_ST_* test where *Q_ST_* is compared to a simulated distribution of *F_ST_*, this test still does not fully account for the demographic relationships among the populations, because it is reduced to a single summary of the data, the *F_ST_*. Further, the *Q_ST_* – *F_ST_* test may also give misleading results because each trait is considered in isolation, and the signature of adaptive trait differentiation can be the result of selection at another trait if they are genetically correlated (e.g. Chenoweth & Blows, 2008). Ovaskainen et *al*. (2011) proposed an alternative method, which uses a statistically more powerful null hypothesis than the *Q_ST_* – *F_ST_* tests: it accounts for drift distances among all populations (i.e. the coancestry matrix) to derive the null expectation of trait divergence. So far, however, only a handful of applications of this method have been published (e.g. De Kort et *al*., 2016).

Here, we assess adaptive divergence at size, growth and phenology traits in silver fir (*Abies alba* Mill.), an ecologically and economically important European forest tree species. Indeed, recent dendrological (Vitali et *al*., 2017) and genecological (Frank et *al*., 2017a) evidence suggests silver fir to potentially be better adapted to future climatic conditions, notably to drought, than the other dominant conifer of European forests, Norway spruce (*Picea abies* Karst.), which often co-occurs with silver fir. In this study, we concentrate on populations from ecologically diverse sites across Switzerland, including populations from core ranges (Pre-Alps, Central Plateau and Jura Mountains), but also marginal populations isolated by high mountain ranges. These alpine populations encompass the wet outer Alpine chain to the dry Central Alps across diverse soil types (Ellenberg, 1988). Silver fir populations in Switzerland are rather young, palynological and dynamic vegetation modeling suggest that the Swiss Alps were colonized by silver fir from south to north after the Last Glacial Maximum. The species most likely reached the southern slopes of the Alps between 10 and 9 kyr BP and the northern slopes between 8 and 5 kyr BP (Van der Knaap *et al*., 2005; Liepelt *et al*., 2009; Ruosch *et al*., 2016). Range-wide patterns of chloroplast and mitochondrial DNA variation (Liepelt et *al*., 2002) and isozyme data (Burga & Hussendörfer, 2001) from extant silver fir populations suggest that the Swiss Alps were colonized from a single ancestral refugial population situated in the Central and/or Northern Apennines, even though the potential contribution of eastern refugial populations cannot be excluded.

The potential for adaption can be the greatest in isolated populations and poor environmental conditions, however, gene flow from the core areas can easily counteract selection (Lenormand, 2002). Many other wind-pollinated conifer species have high levels of pollen-mediated gene flow (Petit & Hampe, 2006; Kremer et *al*., 2012). Thus, regardless of the heterogeneous environment across Switzerland, little adaptive trait differentiation is expected among the studied populations. Indeed, results of a large-scale common-garden experiment involving seedlings from 90 silver fir populations across Switzerland show that levels of population divergence are rather low, particularly when compared to those of Norway spruce from the same area (Frank et *al*., 2017b). For example, population divergence(*Q_ST_*) in seedling height for silver fir was 0.22 and in phenological traits ≤ 0.11. Nevertheless, Frank et *al*. (2017b) also found that *Q_ST_* values of these traits are much higher than population differentiation at isozyme loci (*F_ST_* = 0.034) in a similar set of populations (Finkeldey et *al*., 2000) and speculated that population divergence might reflect adaptation to local environmental conditions. A further evidence for possible adaptation was that population differentiation in seedling height showed a regional pattern, i.e. seedlings from the Central Plateau were taller than those from the Alps, and environmental variables explained a large proportion of the trait differentiation (49%) (Frank et *al*., 2017b).

Here, we revisit the phenotypic data obtained by Frank *et al*. (2017b) for a subset of 19 populations, complement it with modern genetic data, and assess to what extent the observed trait differentiation is due to selection. The goal of this study is three-fold. First, we test for adaptive trait divergence using the multi-trait test developed by Ovaskainen *et al*. (2011), thus we account for the full demography in devising the null expectation for trait divergence. Since, we have data from a relatively large number of populations and from a highly heterogeneous environment separated by potential barriers to gene flow, accounting for the likely complex and hierarchical population structure is essential. Further, the studied set of size, growth and phenology traits most likely do not evolve independently, thus the multi-trait approach is highly desirable to identify any correlated responses to selection. Second, we develop a set of downscaled historical climate variables and adopt soil data from Frank et *al*. (2017b) to test which one of them drives the adaptive divergence patterns using the methodology described in Karhunen et *al*. (2014). Third, we wish to advocate the methodology developed herein to leverage information that can be obtained from a common-garden trial for testing hypothesis about adaptive evolution. In particular, we contrast our results with the simple *Q_ST_* – *F_ST_*, and propose two methodological improvements to evaluate the signature of selection in each population and a more powerful version of the H-test proposed by Karhunen et *al*. (2014).

## Material and Methods

### Study sites and sampling

In 2009, 90 silver fir stands were selected across Switzerland in the framework of a genecological study (Frank et *al*., 2017b). From each population, three mother trees were selected from an area with a relief as uniform as possible and with at least 100m distance between them. From this study, we selected a subset of 19 presumably allochthonous populations that represented the main geographic regions of Switzerland (Figure 1A), and re-visited them in 2013 and 2016. In spring 2013, needles were sampled from nine to ten adult trees, including the three mother trees, if they were found. In spring 2016, needles were sampled from an additional ten to twelve trees, always respecting a minimum distance of 100m between the newly or previously sampled trees. In total, 387 adult trees were used in this study, with at least 20 individuals per population. Needles were stored in plastic bags for a maximum of one week at 5°*C* before being freeze-dried and prepared for DNA extraction.

**Figure 1.**
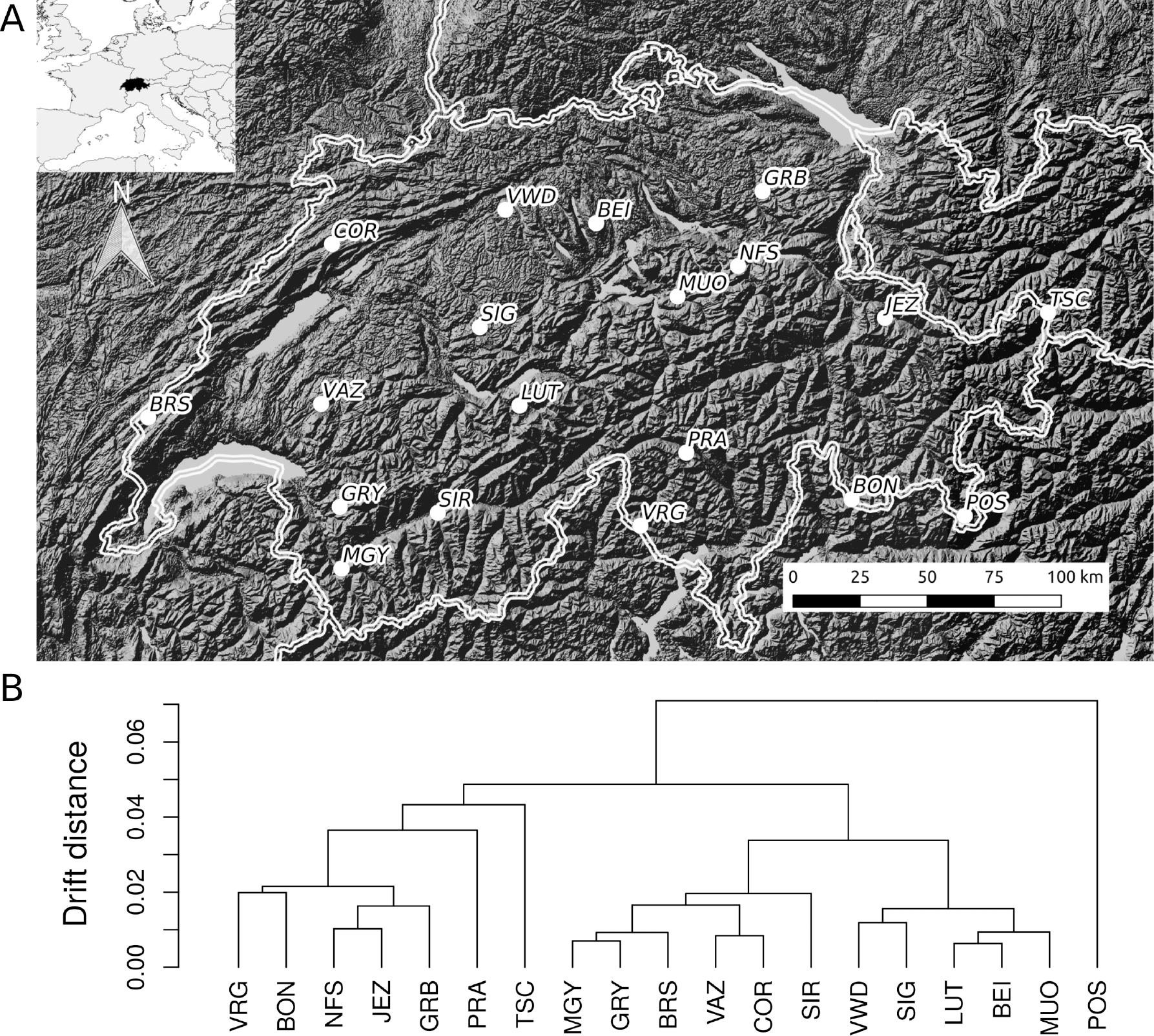
A: Geographic location of the sampling sites. B: Drift distances between populations as estimated with the admixture F-model (AFM). Coancestry between populations is the mean of the posterior means from 10 independent Markov chains. Distances were calculated from the coancestry matrix to draw the dendrogram.

### Genetic data

All adult trees were genotyped at 374 single-nucleotide polymorphism (SNP) loci originating from three different sources. First, we used SNPs discovered from 13 *A. alba* trees originating from Mont Ventoux (France) and the Black Forest (District Oberharmersbach, Germany). Transcriptomes were sequenced on an Illumina HiSeq2000 at IGA Technology Services (Udine, Italy). Transcriptome assembly and SNP detection was carried out with an in-house pipeline at IGA; see details in Roschanski *et al*. (2013). Previous SNP genotyping using KASP assays manufactured by LGC Genomics (Middlesex, United Kingdom) in French populations yielded 267 polymorphic loci situated in 175 different candidate genes (Roschanski et *al*., 2016). In this study, 220 of these loci were successfully genotyped using the same KASP assays. Second, we selected 110 new putatively neutral SNPs from the transcriptome assembly based on their Tajima’s D being between 2 and −2 and dN/dS between 0.9 and 1.1, low LD with the existing 224 SNPs (r2 < 0.1 and p-values > 0.05). ESTScan version 3.0.3 (Iseli et *al*., 1999) for *Arabidopsis thaliana* was used to predict coding DNA sequences for each contig from the transcriptome (Roschanski *et al*., 2013), then the SNPs detected by IGA were used to create pseudo-contigs on which the neutrality statistics were calculated. dN/dS was calculated with ‘polydNdS’^1^ based on (Thornton, 2003), and Tajima’s D with VariScan v2.0.3 (Vilella *et al*., 2005). Although all 110 SNPs passed initial in silico design tests, only 25 of these SNPs were successfully genotyped using KASP technology, most likely because the primer sequences were not specific enough (results not shown). Third, we selected 149 SNPs from Mosca *et al*. (2012), who discovered SNPs by resequencing 800 amplicons using PCR primer pairs from *Pinus taeda* in 12 *Abies alba* trees, and genotyped on a Golden Gate platform (Illumina, San Diego, CA, USA). We selected SNPs from this study that had less than 5% missing data. One hundred and twenty-nine of these SNPs were successfully genotyped with KASP. Both DNA extraction and all genotyping was performed using the all-inclusive service from LGC Genomics (Middlesex, United Kingdom).

### Common garden experiment

We used published seedling phenotype data from the descendants of three mother trees from each of the 19 populations (Frank *et al*., 2017b), while correcting inconsistencies and outliers based on measurements taken in subsequent years (unpublished data of Andrea Kupferschmid). Here, we briefly summarize the protocol of the experiment relevant for silver fir. In April 2010, from each mother tree approximately 2000 seeds (subsequently called family) were sown directly into nursery beds and grown for two years at the Swiss Federal Research Institute WSL in Birmensdorf, Switzerland (47°21′42″*N*, 8°27′22″*E*, 550 m a.s.l.). Families and populations were not replicated or randomized in the nursery. In spring 2012, at least 12 randomly selected viable seedlings per family with present terminal buds were transplanted to the open experimental field site. The field site was located at Brunnersberg, a former pasture on a south facing slope (20-24% incline) in the Swiss Jura Mountains (47°19′35″*N*, 7°36′42"*E*, 1090 m a.s.l.). Seedlings were planted at 30 × 40 cm spacing in 10 blocks. Provenances, families and seedlings were randomized across blocks. Mortality during 2012 in the field was minor. Seedling height and diameter were measured in the autumn of 2012 and used in the analysis as a covariate to account for possible growth differences in the nursery.

Phenotypic measurements used herein were performed during the growing season of 2013. Measurements of seedling Height (variable names capitalized hereafter) and Terminal and Lateral Bud Break started on 7 May 2013, when the first seedlings started to break buds, i.e. the membrane below bud scales was broken and the first green needles became visible. Observations of bud break dates continued twice a week until Growth Cessation, defined as the date when 95% of terminal leader height growth was achieved. Five to 17 height measures were recorded per seedling from which a Maximum Growth Rate was calculated as the first derivative of the growth curve (for simplicity, subsequently called Growth Rate). Growth Duration was calculated as time from Terminal Bud Break to Growth Cessation. Final seedling Height from ground surface to the uppermost bud base and stem Diameter at 2cm above ground surface were measured in autumn 2013. Finally, we also calculated the ratio Height/Diameter to test for trait divergence in growth allometry. For clarity, we call Height, Diameter, Height/Diameter size traits, Growth Rate and Duration growth traits, and Terminal/Lateral Bud Break and Growth Cessation (equivalent to bud set) phenology traits. All traits were normally distributed and correlated with one another to a varying extent (Figure S1).

### Environmental data

We used soil characteristics and downscaled historical climatic data to characterize environmental conditions in each population. Soil and geographic variables were adopted from Frank et *al*. (2017b), who established soil pits within a few meters of one of the mother trees (see list of variables used in Table S1). In order to obtain the closest representation of the climate of the period when the current populations were established, we used data from 1 January 1901 to 31 December 1978. The choice of this period was justified by two facts: (i) no observation-based climate data go back further in time, and (ii) starting from approximately 1980, the temperature time series are overwhelmed by the effect of global warming (Harris et *al*., 2014). We used statistical downscaling using the delta method (Hay et *al*., 2000) to obtain 1 km grid scale mean monthly temperature and precipitation fields for this period. The reference climatic data set was the 0.5° resolution CRU TS v. 4.01 data (20 September 2017 release, Harris et *al*. (2014)) available for the 1 January 1901 - 31 December 2016 period, while the downscaling was based on the overlapping period of 1 January 1979 - 31 December 2016 with the 1 km resolution CHELSA data (Karger et *al*., 2017).

We calculated several bioclimatic indices to characterize the environment for each site, including the 19 bioclimatic variables (Booth et *al*., 2014) implemented in the R package dismo (Hijmans et *al*., 2017), two potential evapotranspiration (PET) indices and four standardized precipitation - evapotranspiration index (SPEI) variables using the R package SPEI (Beguería & Vicente-Serrano, 2017), and two indicators of late frost (Table 1). Soil available water capacity (AWC) was obtained at a 250 m resolution from the Soilgrids data base (Hengl et *al*., 2017). Using temperature, precipitation and AWC, we calculated the self-calibrated Palmer’s drought severity index or scPDSI (Wells et *al*., 2004), and summarized to measures of drought severity and frequency across the full time series (Table 1). All climatic variables were considered as raw values or as deviations from the common garden environment in Brunnersberg (based on the CHELSA data for the period of 1 January 1979 - 31 December 2013). However, the two ways of calculating the climate led to the same conclusions (results not shown), so we present results with the raw variables only for ease of interpretation. To visualize the environmental differences between sites, we conducted a Principal Component Analysis (PCA) of all environmental variables of Tables 1 and S1, scaled and centered to make them comparable. The first PCA explained 32.6% of the variance and was dominated by temperature variables, and the second PCA axis explained 18.8% of the variance and was dominated by precipitation variables (Figure S2).

**Table 1:**
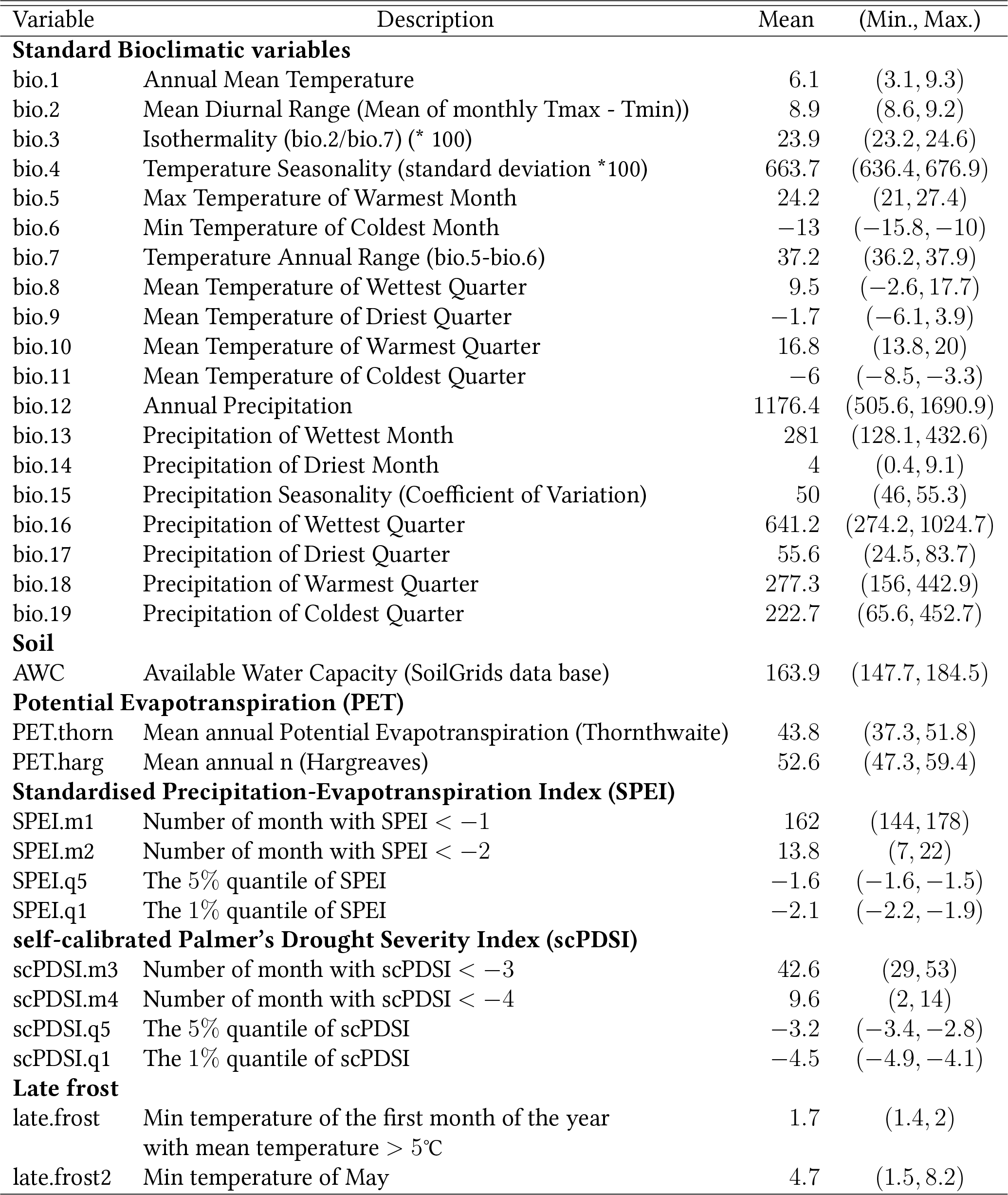
Climatic variables developed in this study using statistical downscaling from CHELSA (Karger et *al*., 2017) and the CRU TS v. 4.01 data (Harris et *al*., 2014). Monthly mean, minimum and maximum temperature and total precipitation were used to calculate the bioclimatic, PET, SPEI and frost variables. Soil AWC was extracted from the SoilGrids data base (Hengl et *al*., 2017), and used for the calculation of the scPDSI variables.

| Variable | Description | Mean | (Min., Max.) |
| --- | --- | --- | --- |
| **Standard Bioclimatic variables** |  |  |  |
| bio.1 | Annual Mean Temperature | 6.1 | (3.1, 9.3) |
| bio.2 | Mean Diurnal Range (Mean of monthly Tmax - Tmin)) | 8.9 | (8.6, 9.2) |
| bio.3 | Isothermality (bio.2/bio.7) (* 100) | 23.9 | (23.2, 24.6) |
| bio.4 | Temperature Seasonality (standard deviation *100) | 663.7 | (636.4, 676.9) |
| bio.5 | Max Temperature of Warmest Month | 24.2 | (21, 27.4) |
| bio.6 | Min Temperature of Coldest Month | -13 | (-15.8, -10) |
| bio.7 | Temperature Annual Range (bio.5-bio.6) | 37.2 | (36.2, 37.9) |
| bio.8 | Mean Temperature of Wettest Quarter | 9.5 | (-2.6, 17.7) |
| bio.9 | Mean Temperature of Driest Quarter | -1.7 | (-6.1, 3.9) |
| bio.10 | Mean Temperature of Warmest Quarter | 16.8 | (13.8, 20) |
| bio.11 | Mean Temperature of Coldest Quarter | -6 | (-8.5, -3.3) |
| bio.12 | Annual Precipitation | 1176.4 | (505.6, 1690.9) |
| bio.13 | Precipitation of Wettest Month | 281 | (128.1, 432.6) |
| bio.14 | Precipitation of Driest Month | 4 | (0.4, 9.1) |
| bio.15 | Precipitation Seasonality (Coefficient of Variation) | 50 | (46, 55.3) |
| bio.16 | Precipitation of Wettest Quarter | 641.2 | (274.2, 1024.7) |
| bio.17 | Precipitation of Driest Quarter | 55.6 | (24.5, 83.7) |
| bio.18 | Precipitation of Warmest Quarter | 277.3 | (156, 442.9) |
| bio.19 | Precipitation of Coldest Quarter | 222.7 | (65.6, 452.7) |
| **Soil** |  |  |  |
| AWC | Available Water Capacity (SoilGrids data base) | 163.9 | (147.7, 184.5) |
| **Potential Evapotranspiration (PET)** |  |  |  |
| PET.thorn | Mean annual Potential Evapotranspiration (Thornthwaite) | 43.8 | (37.3, 51.8) |
| PET.harg | Mean annual n (Hargreaves) | 52.6 | (47.3, 59.4) |
| **Standardised Precipitation-Evapotranspiration Index (SPEI)** |  |  |  |
| SPEI.m1 | Number of month with SPEI < -1 | 162 | (144, 178) |
| SPEI.m2 | Number of month with SPEI < -2 | 13.8 | (7, 22) |
| SPEI.q5 | The 5% quantile of SPEI | -1.6 | (-1.6, -1.5) |
| SPEI.q1 | The 1% quantile of SPEI | -2.1 | (-2.2, -1.9) |
| **self-calibrated Palmer's Drought Severity Index (scPDSI)** |  |  |  |
| scPDSI.m3 | Number of month with scPDSI < -3 | 42.6 | (29, 53) |
| scPDSI.m4 | Number of month with scPDSI < -4 | 9.6 | (2, 14) |
| scPDSI.q5 | The 5% quantile of scPDSI | -3.2 | (-3.4, -2.8) |
| scPDSI.q1 | The 1% quantile of scPDSI | -4.5 | (-4.9, -4.1) |
| **Late frost** |  |  |  |
| late.frost | Min temperature of the first month of the year with mean temperature > 5°C | 1.7 | (1.4, 2) |
| late.frost2 | Min temperature of May | 4.7 | (1.5, 8.2) |

### Statistical analysis

We used both single-trait and multi-trait approaches to test if the phenotypic divergence between populations is due to selection. First, we used a single-trait *Q_ST_* – *F_ST_* test using the bootstrap procedure described in Whitlock & Guillaume (2009) implemented in the R package QstFstComp (Gilbert & Whitlock, 2015)^2^. We considered a one-tailed test, because we were interested in testing for adaptive divergence only, thus *Q_ST_* being significantly greater than *F_ST_*. Second, we used single- and multi-trait versions of the test of adaptive divergence (S-test) developed by Ovaskainen et *al*. (2011) and implemented in the R package driftsel^3^. We estimated the coancestry matrix between all pairs of populations from variation in SNP allele frequencies based on the admixture F-model (AFM) using a Metropolis-Hastings algorithm implemented in the R package RAFM^4^ (Karhunen & Ovaskainen, 2012). Thus, we assumed that contemporary populations evolved from a single ancestral population, which is likely a reasonable assumption for silver fir in Switzerland (Liepelt et *al*., 2002; Burga & Hussendörfer, 2001). Ten independent Markov chains of the AFM model reached slightly different local optima using a burn-in of 20,000 iterations followed by 10,000 iterations for estimation with a thinning interval of ten. Running longer chains (up to 50,000 iterations tested, results not shown) did not improve convergence to a common optimum, so we took the mean of the posterior means across the ten chains. Further, to gain more confidence in the estimated coancestry matrix, we compared the posterior mean coancestry matrix against that estimated using the Bayesian clustering algorithm implemented in the software Structure v.2.3.4 Pritchard et *al*. (2000) (see Supporting Information for more details). Third, we used the H-test, developed by Karhunen et *al*. (2014), and a standardized version of it, subsequently referred to *H*\*, to identify the potential environmental drivers of local adaptation. In the following paragraphs, we briefly summarize the rationale of the *S*-test, and its population-wise version, proposed herein, and the rationale of the proposed *H*\* statistic.

Following the notation of Ovaskainen et *al*. (2011), if we consider only additive effects, the vector of additive values of all traits for individual *i* is **a***_i_*, and the matrix of additive vectors for all individuals is **A** = (**a***_i_*)*_i_*. Then, the mean additive value of population *X*, 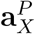, can be obtained as the mean of the additive values of all individuals in population X, and the matrix of additive vectors for all populations is **A**^*P*^ = 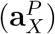*_X_*. When populations are derived from a common ancestral population and the trait values are normally distributed, under drift, the matrix of population-level effects, **A***^P^*, is expected to follow the multivariate normal distribution as

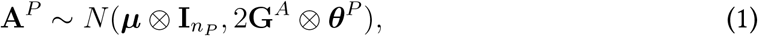

where ***μ*** is the vector of expected additive trait means determined by the allele frequencies in the ancestral population, *n_P_* is the number of populations and **I***_np_* is an *n_P_ × n_P_* identity matrix, ***G**^A^* is the ancestral variance-covariance matrix, ***θ**^P^* is the population-to-population coancestry matrix, and ⊗ is the Kroenecker product. ***θ**^P^* can be estimated assuming the admixture F-model, while ***μ***, **A***^p^*, and ***G**^A^* can be co-estimated using the Bayesian mixed-effects animal model accounting for the family structure of the common garden (i.e. the pedigree) and ***θ**^P^*(Ovaskainen et *al*., 2011). Then, the additive genetic variance-covariance matrix of the contemporary populations assuming no selection, **G**, can be estimated as **G** = 2**G***^A^*(1 – *θ^S^*), where *θ^S^* is the mean within-population (or self) coancestry of all populations, thus the drift distance of the contemporary populations from the ancestral population. Here, we estimated the heritability of traits and the genetic correlation between trait pairs as the proportion of the observed phenotypic variance and covariances that are additive (i.e. using **G**) (Falconer & Mackay, 1996). The evidence for selection can be summarized using the S-statistic calculated as the Mahalanobis distance between **A**^P^ and the distribution of equation 1. *S* = 0.5 indicates consistency with neutrality, S = 0 implies a match with purifying, and S = 1 with diversifying selection. Thus, S measures the overall signature of selection across all populations. In this study, we also test if any particular populations show a signature of selection. We propose that adaptive trait divergence at population X may be concluded if the 95% of the posterior distribution of 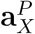 is outside of the neutral envelop defined as ***μ*** ± 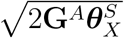. The neutral envelop can be uni- or multi-variate depending if one or multiple traits are studied. Our motivation for this population-wise evaluation of divergence is that a single *S*-statistic cannot distinguish between the two scenarios of many populations that are slightly diverged or a single/few populations that are diverged to a great extent.

The *H*-statistics measure if the distance between the populations’ mean additive trait values is more similar to the environmental distances than expected based on drift (Karhunen *et al*., 2014). The original H-test is based on the Mantel test statistic, i.e. the product moment between the distance matrices of environment and traits. As the covariance is not only influenced by the correlation between the matrices but also by the absolute values of trait differences, the original H-test may yield false positive results. In particular, in cases with strong evidence for selection, the *H* statistic can be high (i.e. close to 1) even when selection is uncorrelated with the tested environmental driver. For this reason, we propose the use of the *H*\* statistic, which is the Pearson or standardized Mantel statistic, thus the *H*\*-test can be viewed as a standardized version of the H-test. We note that the superiority of a standardized Mantel statistic has already been pointed out in the context of spatial distance matrices (Legendre & Fortin, 2010). The population-wise divergence test and the *H*\*-test are now implemented in the most recent version of the R package driftsel^5^.

## Results

### Population structure and drift distances

*F_ST_* using the Whitlock & Guillaume (2009) approach was 0.0056 (95% confidence interval: 0.0051, 0.0061). The drift distances estimated with AFM indicated the presence of two main clusters that correspond to eastern and western populations (Figure 1B). The population POS did not belong to either of these groups, which is plausible given that it is situated south of a high mountain pass (the Bernina pass) close to the Italian border (Figure 1A). The coancestry matrix can also be used to obtain a coancestry-based estimate of *F_ST_*, which was 0.0184 (95% credible interval: 0.0167, 0.0202) for the 19 populations. The comparison between AFM and Structure revealed similar results (see Supporting Information for more details).

### Single-trait analysis

Single-trait tests of adaptive divergence in terms of *Q_ST_* – *F_ST_* comparisons using the Whitlock & Guillaume (2009) approach and in terms of the *S* statistic from the Ovaskainen *et al*. (2011) method gave similar results (Table 2). Using either of the methods, the strongest signature of adaptive divergence was observed for seedling Height followed by four other traits with a strong evidence of selection; in decreasing order of evidence based on the *S* statistic: Lateral Bud Break, Terminal Bud Break, Diameter, and Growth Duration. The remaining three traits showed no or a weak signal of unusual trait differentiation using either of the approaches. For Growth Cessation and Height/Diameter the *Q_ST_* – *F_ST_* test was marginally significant, while the S statistic was close to 0.5, consist with neutrality. For Growth Rate, there was no evidence of selection using either approach.

**Table 2:** Summary of single-trait tests of adaptive divergence. *Q_ST_*, *Q_ST_* – *F_ST_* and p-value correspond to the Whitlock & Guillaume (2009) approach, *S* is the summary of evidence of selection using the Ovaskainen et *al*. (2011) method, *H* indicates the role of the environment in adaptation using the Karhunen et *al*. (2014) test, and *H*\* is a standardized version of *H* proposed in the study. Both *H* and *H*\* were calculated with all environmental variables listed in Tables 1 and S1. Values of *S*, *H*, and *H*\* close to 1 indicate high evidence of divergent selection.

| Trait | $Q_{ST}$ | $Q_{ST} - F_{ST}$ | p-value | S statistic | H statistic | $H^*$ statistic |
| --- | --- | --- | --- | --- | --- | --- |
| Diameter | 0.082 | 0.076 | 0.032 | 0.953 | 0.971 | 0.883 |
| Height | 0.268 | 0.262 | 0.001 | 1.000 | 1.000 | 0.930 |
| Height/Diameter | 0.245 | 0.239 | 0.045 | 0.864 | 0.449 | 0.606 |
| Growth Rate | 0.060 | 0.054 | 0.113 | 0.768 | 0.828 | 0.703 |
| Growth Duration | 0.275 | 0.269 | 0.031 | 0.944 | 0.905 | 0.473 |
| Terminal Bud Break | 0.212 | 0.206 | 0.020 | 0.958 | 0.880 | 0.534 |
| Lateral Bud Break | 0.138 | 0.132 | 0.016 | 0.964 | 0.885 | 0.480 |
| Growth Cessation | 0.436 | 0.430 | 0.045 | 0.551 | 0.425 | 0.355 |

The population-wise *S*-tests revealed a signature of selection in several populations. Figure 2 shows, for each trait, how much each population diverged from the ancestral mean and if this divergence is more than expected by drift. These population-level estimates confirmed that Height diverged the most among populations (Figure 2B): eight out of 19 populations deviated from the neutral expectation with at least 95% posterior probability. All these eight populations evolved towards a higher mean height and no populations have been selected for reduced height. Although the deviation from the neutral expectation was not as strong for Diameter as for Height, the first seven tallest populations also diverged towards the largest Diameter (Figure 2A), suggesting co-evolution between Height and Diameter, and explaining the lack of unusual differentiation at Height/Diameter (Figure 2C). Evidence for trait divergence was also observed for a single population, SIR, for Growth Duration (Figure 2E) and Lateral Bud Break (Figure 2H). The SIR population evolved towards earlier bud break and longer growth duration than expected by drift alone. At the opposite end of the trait space, population VRG evolved towards late Terminal Bud Break and shorter Growth Duration, yet the evidence for divergence was not very strong in this single-trait test.

**Figure 2.**
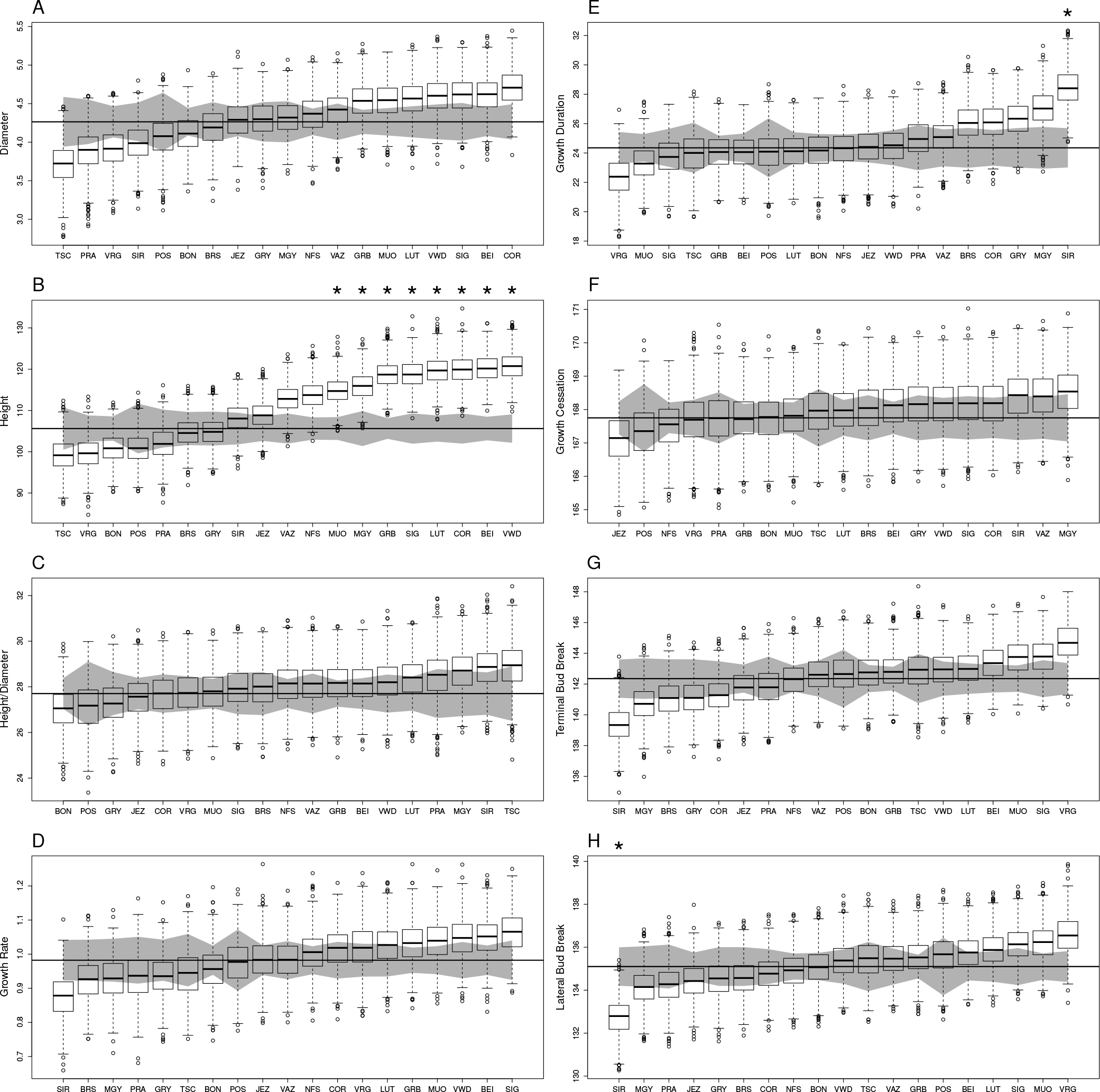
Summary of the single-trait test of adaptive divergence using the Ovaskainen et *al*. (2011) method for the eight studied traits shown across panels A to H. Each panel shows the estimated ancestral additive mean trait value (horizontal line), the amount of trait divergence from this mean that is expected based on drift (gray envelop), and the estimated posterior distribution of the additive trait values for each population (boxes). Stars above the boxes indicate the signature of selection at the particular population. Population order is different on each panel because populations are ordered according to their additive trait values.

### Multi-trait analysis

We performed only two-trait analyses due to convergence problems with three or more traits (results not shown). We considered only one of the bud break traits in this pairwise analysis, Terminal Bud Break, because the two traits were very strongly correlated. The heritability values for a given trait were slightly different depending on the particular pairwise analysis because **G^A^** is different, and also *θ*^**P**^ is updated in the Bayesian animal model (Table S2). For most trait pairs the genetic correlations were small or negligible (Table S2) and agreed well with phenotypic correlations (Figure 3A). The highest phenotypic correlation (0.64) was observed between two size traits, Height and Diameter, yet their genetic correlation was half as much (0.32, Table S1). Ten populations showed evidence for adaptive co-divergence in Height and Diameter (Figure 4). In contrast, some unusually high genetic correlations (i.e. higher than the phenotypic equivalents) were observed between traits of different kind: between growth and phenology (Growth Duration and Terminal Bud Break, Growth Rate and Terminal Bud Break), and between size and phenology (Height/Diameter and Growth Cessation; Figure 3A). The S-statistics was high only for the first two trait pairs (Figure 3B), suggesting a correlated evolution at the three traits (Figure 5). In particular, two populations reached an important level of divergence in the opposite directions. These patters can be interpreted as contrasting life history strategies. SIR showed adaptive divergence for all three trait combinations, while MGY for Growth Duration vs Growth Rate (Figure 5). Thus, we can say that SIR and MGY follow a “start early and grow slow” strategy, i.e. they burst buds early and then grow for a long time at a low rate. At the other end of trait space, population VRG follows a or “start late and grow fast” strategy, i.e. bursts buds late, but then grows fast for a short period of time (Figure 5).

**Figure 3.**
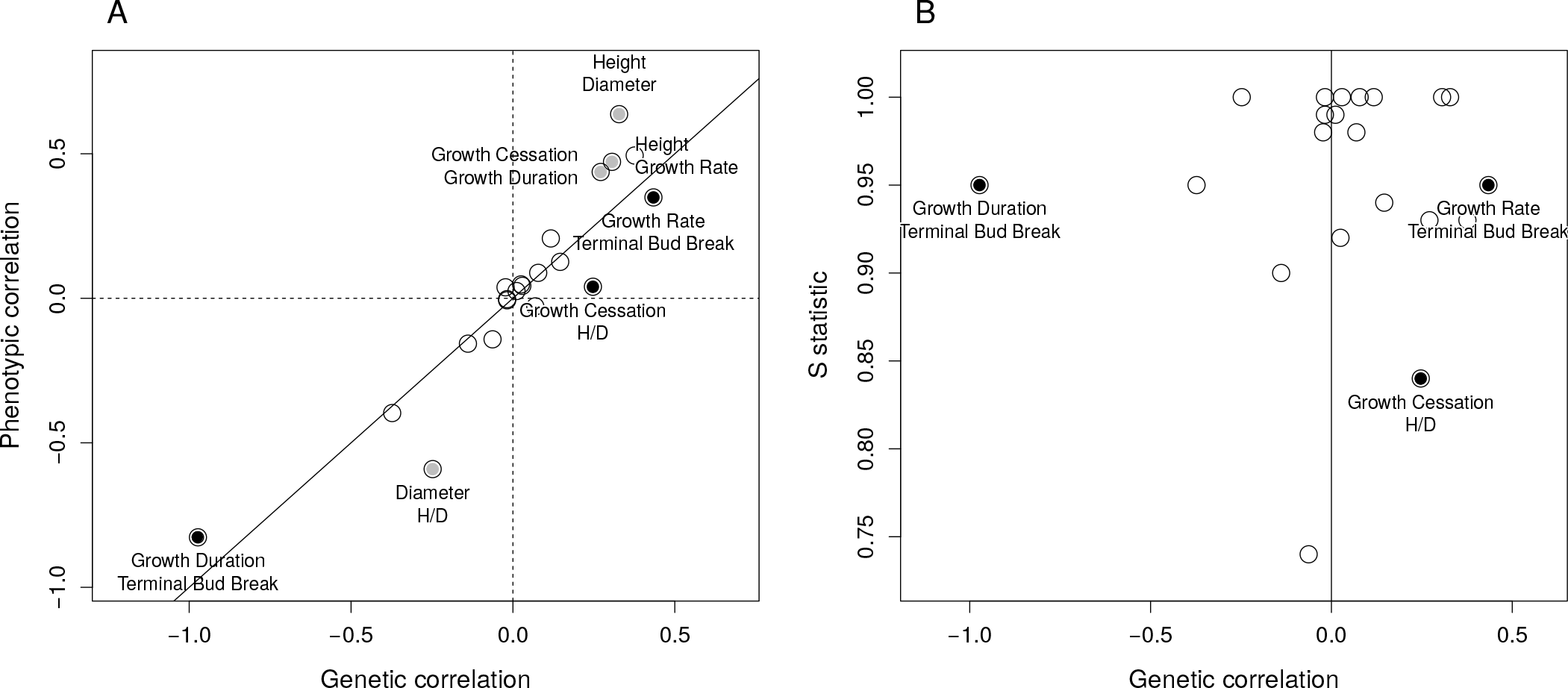
Summary of the two-trait analysis of adaptive divergence using the Ovaskainen et *al*. (2011) method. Panel A shows the phenotypic (*r_p_*) and genetic (*r_g_*) correlations between trait pairs (see main text for formula). The top three trait pairs with *r_p_* ≥ *r_g_* are highlighted in gray, and those with *r_g_* ≥ *r_p_* in black. Panel B shows the S-statistic for the two-trait selection test as a function of *r_g_*. The three trait pairs with *r_g_* > *r_p_* highlighted in panel A are also highlighted here.

**Figure 4.**
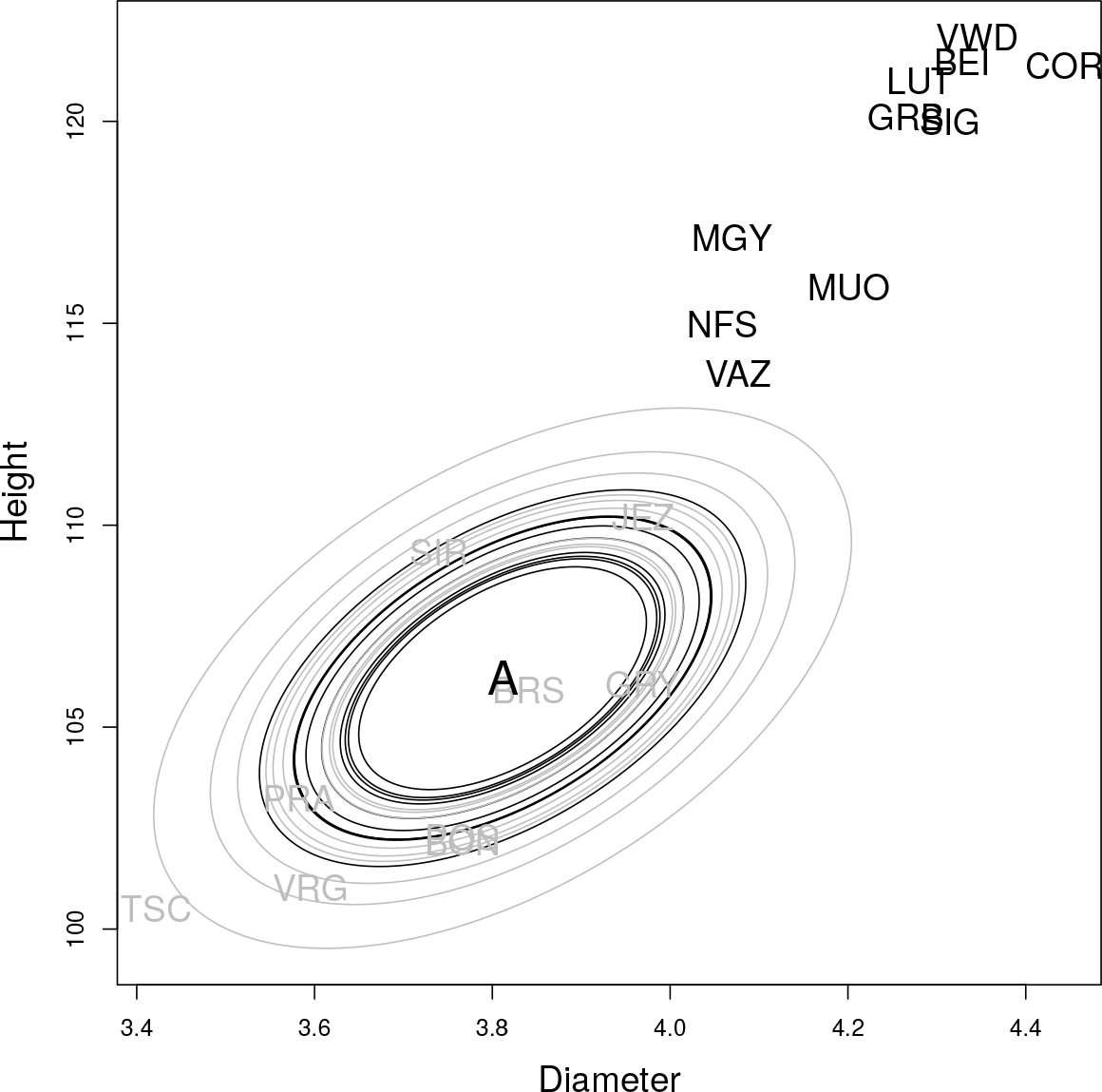
Correlated adaptive divergence in Height and Diameter using the Ovaskainen et *al*. (2011) method. *A* is the estimated ancestral additive mean trait value. Ellipses represent the median amount of trait divergence that is expected based on drift for each population. Population codes (3 letters) represent the median of the posterior distribution of the additive trait values for each population. Ellipses and population codes with evidence of trait divergence are shown in black, while populations that do not deviate from what is expected based on drift are in gray.

**Figure 5.**
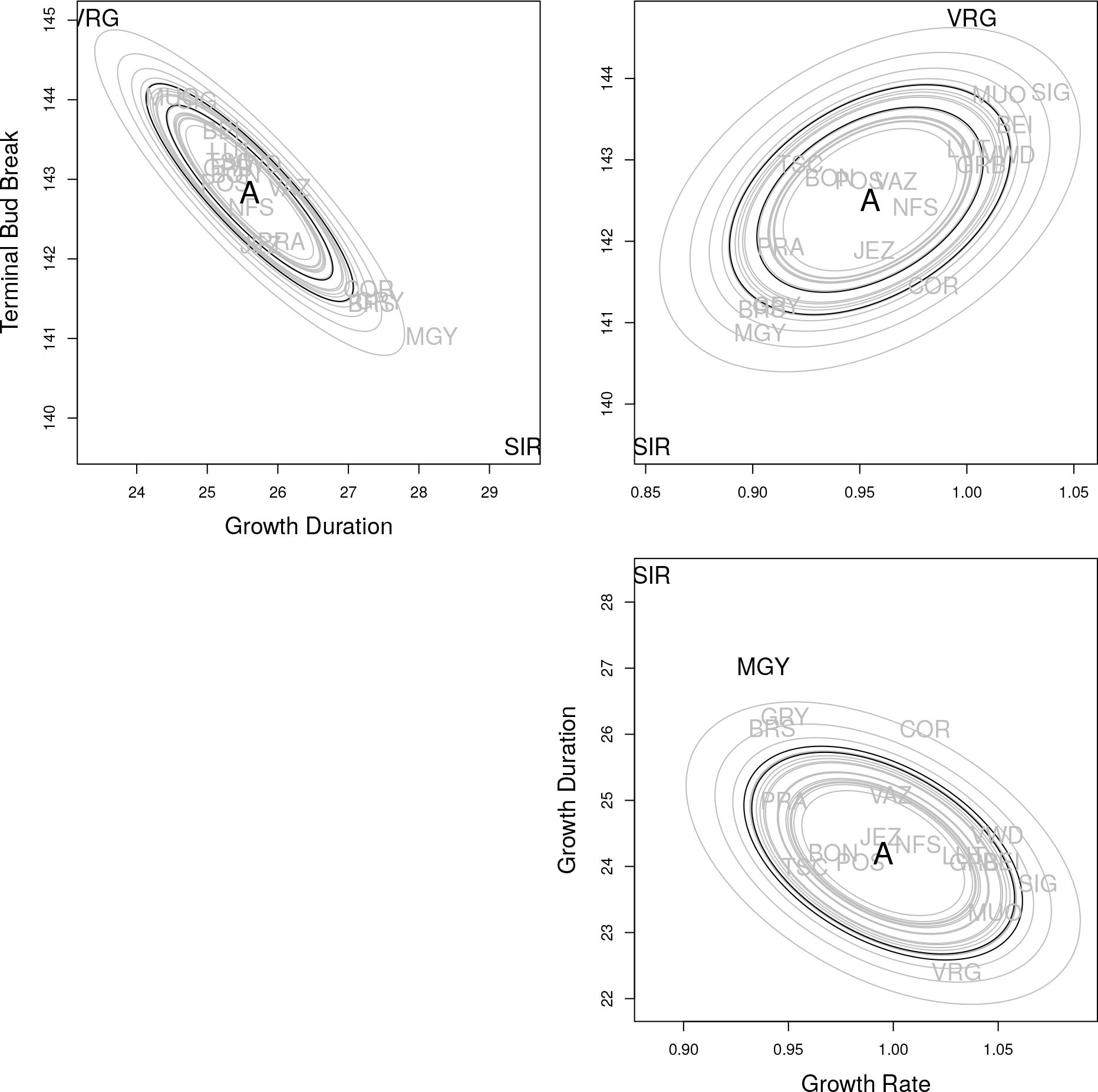
Correlated adaptive divergence at Terminal Bud Break, Growth Duration and Growth Rate using the Ovaskainen *et al*. (2011) method. Each panel shows the evidence for unusual trait divergence in a two-trait space. A is the estimated ancestral additive mean trait value. Ellipses represent the median amount of trait divergence that is expected based on drift for each population. Population codes (3 letters) represent the median of the posterior distribution of the additive trait values for each population. Ellipses and population codes with evidence of trait divergence are shown in black, while populations that do not deviate from what is expected based on drift are in gray.

### Environmental drivers

*H* and *H*\* assess how much each environmental variable moves the Mantel statistic away from the neutral expectation (i.e. random association with the environment conditional on the drift distances). First, we tested the effect of the “global” environment, i.e. used all environmental variables listed in Tables 1 and S1 to calculate the environmental distance matrix. We repeated this analysis on the scaled and centered environmental variables, but they led to similar results, so only the analysis using the raw variables is reported. The “global” environment explained trait divergence for Diameter, Height and Growth Rate (Table 2). For traits for which S was close to 0.5, such as Height/Diameter and Growth Cessation, there were no associations with the environmental variables either. Similarly, for Bud Break traits and Growth Duration, *H* was also slightly smaller than *S* and *H*\* was close to 0.5 (Table 2). However, using the “global” environment, we assume that all aspects of the environment are important for adaptation. Further such global environmental distance matrix is dominated by the effects of temperature, which explains the most variation in the environmental data (Figure S2). In reality, it is likely that for each particular trait a particular aspect of the environment matters more than other aspects, so we applied the *H*- and *H*\* -tests for each environmental variable listed in Tables 1 and S1 separately. Since our objective was to prioritize the environmental variables, we did not correct for multiple testing. The *H* - and *H*\*-statistics generally led to similar conclusions concerning the role of each environmental variable, but the *H*-statistic was inflated under a generally strong signal of selection, such as for Height (Figure S3).

The role of the environment in the divergence of Height and Diameter was most important among all traits (Table 2). Additionally, the environmental variables that had an *H*\*-statistic higher than 0.95 for Height included all variables that fulfilled the same criterion for Diameter, suggesting that the same environmental cues matter for the evolution of both traits. Remarkably, all of these environmental variables were related to temperature and drought, but not precipitation per se. The most important environmental variables in the order of decreasing *H*\*-statistic for Height were Thornthwaite’s mean annual evapotranspiration (PET.thorn), mean temperature of the warmest quarter (bio.10), isothermality (bio.3), annual mean temperature (bio.1), available water capacity (AWC), elevation, mean temperature of the coldest quarter (bio.11), maximum temperature of the warmest month (bio.5), minimum temperature of the coldest month (bio.6), mean diurnal range (bio.2), latitude and drought severity measured as the 1% quantile of the self-calibrated Palmer’s Drought Severity Index across the whole time series. To support the interpretation of these results, we performed a PCA of these environmental variables. We found that all populations that diverged towards a larger size in terms of Height and Diameter were situated at the warm, stable and humid side of the environmental space (Figure 6A). Finally, note that the precipitation-related variables (bio.12, bio.13, bio.14, bio.15, bio.18) had the lowest correlation with adaptive divergence for tree size (*H*\*-statistics were below 0.5).

**Figure 6.**
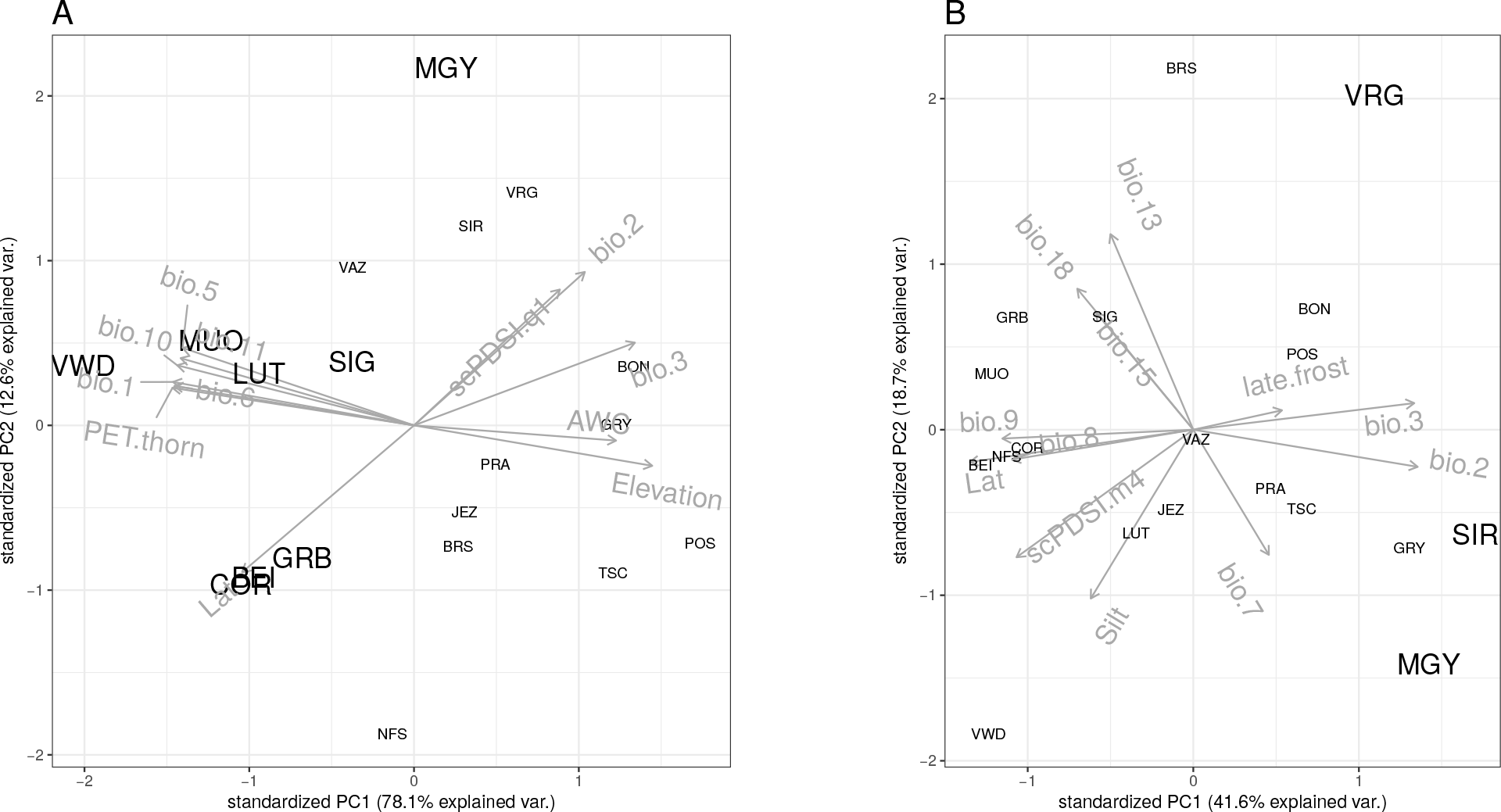
A: The first two principal components of the environmental variables that had a *H*\* -statistics higher than 0.95 for Height and Diameter. Populations that showed a signature of adaptive divergence for Height in the population-wise *S*-test are situated in the “warm” and “stable” left side of the environmental space and are indicated by larger letter sizes for their population codes (i.e. VWD, BEI, COR, LUT, SIG, GRB, MGY, MUO). B: The first two principal components of the environmental variables that had and *H*\*-statistics higher than 0.85 for Growth Rate and Duration, and Terminal and Lateral Bud Break. Populations that showed a signature of adaptive divergence for these traits in the single- or multi-trait population-wise *S*-test are indicated by larger letter sizes for their population codes (i.e. SIR, MGY, VRG). Explications of the environmental axes (in gray) can be found in Table 1.

Divergence at the two Growth traits and the two Bud Break traits also correlated with habitat differences: The most important environmental variables (*H*\*≥0.85) largely overlapped for these traits and were most often related to precipitation (Figure 6B). Mean temperature of the wettest quarter (bio.8), precipitation seasonality (bio.15), precipitation of warmest quarter (bio.18), and mean diurnal range (bio.2) were omnipresent as environmental variables with the highest *H*\*-statistics for the four traits. Again, the common habitat variables that explain adaptation for these traits is not surprising given the evidence for their correlated evolution (Figure 5). The PCA of these environmental variables illustrates that the drivers of the “start early and grow slow” strategy of SIR, and to some extent that of MGY and GRY, are low average precipitation, low of precipitation seasonality, and high temperature amplitude both at the annual and monthly levels (Figure 6B), reflecting the cold winters and hot and dry summers of the Rhône Valley. At the other end of the trait space, the climate of population VRG is characterized by ample winter precipitation, late snow and late frost, and low soil silt content, which could explain the late bud break and short but fast growth of this population (Figure 6B).

## Discussion

In this study, we found clear evidence of local adaptation at size, growth and phenology traits of seedlings originating from 19 silver fir populations from across Switzerland. Thus, our study relies on the common, and often supported (*e.g*. Cornelissen et *al*., 2003; Major et *al*., 2003), assumption that traits measured at seedlings predicts those at adult trees. Tree size (Height and Diameter) has been under strong selection in warm and humid areas of Switzerland (Central Plateau), where populations evolved towards a taller and larger stature (Figure 4 and 6). The role of selection on Height is further supported by the fact that the selection gradient is South-North (northern populations being taller), while the main demographic axis is East-West (Figure 1B). The annual development cycle of temperate trees likely results from an evolutionary trade-off between two opposing forces: maximizing the length of the vegetative season while avoiding late frost and summer drought. This life-history trade-off is particularly crucial in mountainous environments, where the length of the growing season is often limited. In this study, the timing, the rate, and the duration of growth have been under selection that allowed populations to evolve towards two extreme strategies that reflect adaptation to the amount and timing of precipitation (Figure 5 and 6). Populations of the Rhône Valley, especially SIR, but also MGY and GRY (Figure 1A), seem to persevere the conditions of this notoriously dry inner Alpine valley of Switzerland. Here, annual precipitation is low and falls mainly in winter, while high temperatures and drought are common during spring and summer. These populations are adapted to such climatic conditions through early bud break and slow growth during the dry but long vegetative season (Figure 5). We observed the opposite strategy in VRG, and to some extent in MUO. Both populations are situated in high-altitude mountain valleys (Figure 1A), where late snow delays the start of the vegetative season, but then growth is fastest in these populations (Figure 5).

Quantitative genetic theory suggests that fitness-related traits are under strong selective pressure with stabilizing selection reducing genetic variation among individuals (e.g. Mousseau & Roff, 1987). As a result, it is often observed that heritability estimates are lower on morphological than on life-history traits (Visscher et *al*., 2008). In contrast, estimates of heritability from Frank et *al*. (2017b) using 90 Swiss populations of silver fir revealed the highest heritability for phenology and growth, and lowest for size. Further, our estimates of the heritability derived from the ancestral G-matrix also revealed the highest values for the two Bud Break traits, while the lowest for Height (Table S2). We may speculate that height has been under strong selection, thus its additive genetic variance has been reduced. The fitness advantage of being tall for trees are clear: Taller trees have access to more light, thus can have enhanced growth, and high stature facilitates the pollen and grain dispersion (Petit & Hampe, 2006). Further, artificial selection for height may also have affected these populations, further reducing the additive genetic variance. The populations that have been selected for being taller in this study (Figure 2B) are located in the geographic region that are the most easily accessible for foresters due to their low elevation and often flat terrain. Since tree height is a key trait from an economical point of view, and there has been a long history of management in this region (*e.g*. Bürgi & Schuler, 2003), we cannot exclude the possibility that there has been artificial selection in these populations.

Interestingly, two traits, Height/Diameter and Growth Cessation, did not show evidence of adaptive divergence, but they were genetically correlated (0.25), while lacking any phenotypic correlation (0.04, Figure 3, Table S1). The allometric relationship between Height and Diameter has been extensively studied in forest trees because it determines the volume of the harvestable stem and carbon storage, and it is also associated with wood quality such as lignin and cellulose content (e.g. Chave et *al*., 2014). The fact that Growth Cessation did not show a signature of selection may seem surprising given the overwhelming evidence of adaptive clines for bud set (a proxy for growth cessation) in many species, including conifers (Alberto et *al*., 2013). The explanation may lie in the deterministic bud development of the *Abies* species (Cooke et *al*., 2012). They produce terminal buds during the summer at the tip of each leading branch shoot and remain dormant during the following winter. Each bud contains a preformed miniature of the terminal and lateral shoots with all the needles that will grow to branches and photosynthesizing needles, respectively, during the following growing season. Thus, given that the timing of the start (bud burst) and the rate (maximum growth rate and growth duration) of this developmental trait have been under selection (Figure 5), the timing of the end cannot be under selection. The positive genetic and negligible phenotypic correlation between Height/Diameter and Growth Cessation may indicate that the trees can only increase their height in relation to their diameter at the price of growing longer; however, the evolution of this developmental process is extremely constrained due to canalization.

A long-standing hypothesis in evolutionary biology is that traits belonging to the same functional and/or developmental group are genetically more integrated than traits with different functions or developmental origins (Berg, 1960; Pigliucci & Preston, 2004). Several empirical studies found evidence that there is greater genetic and phenotypic character integration within suites of functionally or developmentally related traits than between them, e.g. between and within floral vs. vegetative traits in plants (Waitt & Levin, 1998; Baranzelli *et al*., 2014). In this study, we measured exclusively vegetative traits, but three kinds, related to size, growth, and phenology. Indeed, we found that the genetic correlations tended to be higher than phenotypic correlations between different trait kinds than within (Figure 3A), and in particular, between growth and phenology. These results suggest that at the physiological and molecular level, growth and phenology are strongly linked.

Overall, our methodology proved more informative than classical *Q_ST_* – *F_ST_* comparisons, even though the high number of populations leveraged the power of the classical test and led to similar global conclusions (Table 2). First, we were able to pinpoint which populations diverged from a supposed ancestral trait value (Figure 2). Second, the multi-trait analysis allowed us to identify life-history strategies that are under selection, and third, we were able to identify the selective environment. In particular, we found that the same environmental variables explained best the single-trait adaptive population differentiation at traits that were genetically correlated in the multi-trait analysis, which provides particularly strong evidence for the role of temperature and humidity in the evolution of tree size (Figure 6A) and the role of precipitation seasonality in the timing and mode of growth (Figure 6B).

Recent evidence suggests a global increase in climate-driven forest decline that is primarily driven by a combination of rising temperatures and drought stress, so-called hot-drought, e.g. Allen et *al*. (2010). Such conditions foster detrimental agents such as pests (pathogens, herbivores) or physical disturbances such as storms (e.g. Csilléry et *al*., 2017). For Europe, the most recent IPCC report predicts an increase in temperature, along with a decrease in summer precipitation and an increase in winter precipitation (Stocker et *al*., 2014). Although there is abundant evidence of the past evolution of plants in response to climate, such as past range shifts during glaciations (e.g. Petit et *al*., 2003; Aitken et *al*., 2008), the question is the extent to which adaptive evolution can compensate the effects of rapid climate change. Silver fir may already be in danger in some areas of the Mediterranean, where die-back events have been documented (Cailleret et *al*., 2014), or in south-western Europe, where reduced growth patterns have been reported (Gazol et *al*., 2015). Yet, positive effects of climate warming have also been observed in temperate areas, where warming enhanced growth (Gazol et *al*., 2015). The predicted pace of climate change is much faster than it has been during the time since post-glacial expansion/re-colonization, thus assisted migration can provide a practical solution (Aitken & Bemmels, 2016). Based on our results, populations of the Rhône Valley may provide dry tolerant seed sources for future plantations in other parts of Switzerland.

## Footnotes

1 https://github.com/molpopgen/analysis

2 https://github.com/kjgilbert/QstFstComp

3 https://www.helsinki.fi/en/researchgroups/metapopulation-research-centre/rafm-and-driftsel

4 https://www.helsinki.fi/en/researchgroups/metapopulation-research-centre/rafm-and-driftsel

5 https://github.com/kcsillery/driftsel

Author contributions
KC designed the study and obtained the SNP data with help from AW and FG. KC performed all analysis with advice from OO. CS was involved in the published common garden experiment and provided help with data interpretation. KC wrote the manuscript with help from all other authors.

## Acknowledgements

We thank all members of the ADAPT project that provided the phenotypic data for this study, especially Aline Frank, Caroline Heiri and Peter Brang. Additionally, Andrea Kupferschmid helped with correcting some errors in the phenotypic data thanks to her follow-up project (EADAPT). Part of the sampling was conducted by Catherine Folly. The project was funded by a research grant from the Center for Adaptation to a Changing Environment (ACE) at the ETH Zürich. KC was supported by ACE fellowship while obtaining and analyzing the data, and by a Marie Sklodowska-Curie fellowship (FORGENET), while working on the manuscript. We thank Kristian Ullrich for help with the SNP selection, and Dirk Krager with extracting the climatic data from the data bases. The contribution of several field and lab workers was necessary for this project, in particular, we thank René Graf and Olivier Charlandie.

